# Tip of the iceberg? Three novel TOPLESS interacting effectors of the gall-inducing fungus *Ustilago maydis*

**DOI:** 10.1101/2023.06.12.544640

**Authors:** Mamoona Khan, Simon Uhse, Janos Bindics, Benjamin Kogelmann, Nithya Nagarajan, Kishor D. Ingole, Armin Djamei

## Abstract

- *Ustilago maydis* is a biotrophic pathogen causing smut disease in maize. It secretes a cocktail of effector proteins during its biotrophic stages in the host plant, which target different host proteins. One such class of proteins we identified previously is TOPLESS (TPL) and TOPLESS RELATED (TPR) transcriptional corepressors.
- Here we screen 297 U. *maydis* effector candidates for their ability to interact with maize TPL protein RAMOSA 1 ENHANCER LOCUS 2 Like 2 (RELK2) and their ability to induce auxin signaling and thereby identified three novel TPL /TPR interacting effector proteins (Tip6, Tip7 and Tip8). Two of them, Tip6 and Tip7 contain a classical ethylene-responsive element binding factor-associated amphiphilic repression (EAR) motif and interact with maize TPL protein RELK2 in nuclear compartments, whereas Tip8 lacks known TPL interaction motifs and its overexpression *in* non-host plant leads to cell death indicating recognition of the effector.
- By using structural modeling, we show an interaction of Tip6 and Tip7 with the previously crystallized EAR motif binding domain of RELK2. Furthermore, by infection assays with an octuple deletion mutant of *U. maydis*, we demonstrate a role of Tips in *U. maydis* virulence. Our findings suggest the TOPLESS class of corepressors as a major hub of *U. maydis* effector proteins.

## Introduction

*Ustilago maydis* is a biotrophic plant pathogen that belongs to the family of smut fungi *Ustilaginaceae* (Martínez-Espinoza et al., 2002) and is specialized to infect maize (*Zea mays L*.) and its predecessor teosinte (*Zea mays sub*.*sp. mexicana and sp. parvigluminis*). The fungus can infect all aerial parts of the host plant (stems, leaves, tassels, and ears) and locally induces gall formation (Kamper et al., 2006). *U. maydis* has a dimorphic lifestyle, a yeast-like saprophytic form that can be propagated on artificial growth media, and a biotrophic filamentous growth stage, developing when haploid sporidia of compatible mating type recognize, fuse, and develop a dikaryotic filament (Kahmann and Kamper, 2004). After mating, *U. maydis* filaments perceive from the cuticula of the host plant physical and chemical cues that lead to the induction of the formation of appressoria, penetration structures that allow the fungus to grow and proliferate thereafter intra- and intercellularly. Colonized tissues become hypertrophic and host-derived galls are induced in which *U. maydis* filaments undergo karyogamy forming aggregates in apoplastic cavities and are embedded in a mucilaginous matrix. This is followed by hyphae fragmentation and the formation of melanized, black teliospores giving upon-release plants a smutted appearance (Banuett and Herskowitz, 1996).

To promote its host colonization, *U. maydis* suppresses plant defence responses, modifies the plant metabolism in its favour, and manipulates the plant cell cycle to create the environment for its proliferation (Djamei et al., 2011; Redkar et al., 2015; Navarrete et al., 2022; Saado et al., 2022). All these changes involve massive metabolic and transcriptional reprogramming on the host side (Doehlemann et al., 2008; Lanver et al., 2018). To achieve this host reprogramming *U. maydis* encodes in its genome a large arsenal of 476 predicted secreted protein encoding genes (Lanver et al., 2017), of which many are likely so called effectors, that are released into the biotrophic interphase during host invasion to manipulate the host. Recent studies have shed light that some of these effector proteins are translocated to the host cell and target highly conserved plant pathways including transcriptional co-repressors (Darino et al., 2021; Bindics et al., 2022; Navarrete et al., 2022).

TOPLESS (TPL) is the most prominent co-repressor in plants that interacts with numerous other repressors, transcription factors, and adaptor proteins to modulate signaling pathways including phytohormone auxin-, jasmonate / ethylene-, brassinosteroid-, abscisic acid-as well as salicylic acid-signaling and have been recently associated also with plant immunity besides the well-known involvement in plant development (Causier et al., 2012a; Altmann et al., 2020). TPL and TPL Related Proteins (TRPs) belongs to the GRO/TUP family of co-repressors, which are conserved in all eukaryotes (Causier et al., 2012b). There are five TPL/TPR family proteins in *A. thaliana* (Plant et al., 2021) and four in maize (Liu et al., 2019). TPL was initially identified genetically, based on developmental defects occurring in *A. thaliana* mutants bearing mutations in the *TPL* gene causing an abnormal development of the embryo (Long et al., 2006). Later on, TPL was shown to make a complex with transcriptional repressors of Aux/IAA proteins through interaction with the EAR motif present in Aux/IAA proteins and repress auxin mediated responses by binding to the promoters of AUXIN RESPONSIVE FACTORS (ARFs) (Szemenyei et al., 2008; Dinesh et al., 2016). The EAR motif, defined by the consensus sequence patterns of either LxLxL or DLNxxP, is the most predominant form of transcriptional repression motif so far identified in plants (Kagale and Rozwadowski, 2011). This role of TPL in auxin signaling is conserved in land plants ranging from the moss *Physcomitrella patens* to the angiosperms (Causier et al., 2012b). Similar regulatory modes have been shown to control the induction of genes of the jasmonic acid hormone signaling pathway, which plays a role in response to various abiotic and biotic stresses (Chico et al., 2008). Recently, the crystal structure of the rice and *Arabidopsis* TPL was solved. This work revealed that the N-terminal part of the protein forms a very conserved fold, called the TPL domain (TPD), and that TPL proteins form tetramers (Ke et al., 2015; Martin-Arevalillo et al., 2017). The C-terminus of TPL protein contains several WD40 repeats, which recruit HISTONE DEACETYLASEs (e.g. HDA19) and HISTONE METHYLTRANSFERASEs (Collins et al., 2019).

Recently, we have identified seven TPL-interacting *U. maydis* effector proteins, five of which are genetically clustered (Bindics et al., 2022). These effectors interact either with the C-terminal WD-40 domain or the N-terminal TPD domain of the TPL proteins and turn on mainly jasmonic acid/ethylene signaling (Darino et al., 2021) or auxin signaling (Bindics et al., 2022; Navarrete et al., 2022) respectively. The genetically clustered TPL interacting proteins (Tips) are a family of five effector proteins belonging to the cluster 6A of *U. maydis* and are involved in promoting virulence. They interact with TPL/TPRs of both, maize and dicot model plants and compete with Aux/IAA repressors to induce auxin signaling. Very interestingly, Tip1-5 effectors do not contain any canonical EAR motif in their protein sequences. Whereas another TPL interacting *U. maydis* effector NAKED1 (Nkd1) interferes with maize TPL/TPRs through its EAR motif interaction with N-termini of TPL/TPR proteins. This leads to an upregulation of auxin signaling and subsequent disruption of PAMP-triggered ROS burst. The *U. maydis* knockout strain of Nkd1(Δnkd1) shows a strong virulence defect which is not linked to its interaction with TPL/TPRs but indicates an independent second function. In contrast to this, the ROS-burst suppressive activity of Nkd1 is directly linked with the presence of its EAR-motif and its ability to bind to TPL/TPRs revealing a function of TPL/TPRs in PAMP-triggered immunity (Navarrete et al., 2022). Another *U. maydis* effector JASMONATE/ETHYLENE SIGNALING INDUCER 1 (Jsi1) also interacts with maize TPL/TPRs in an EAR-dependent manner. Interestingly Jsi1 binds the second WD40 domain at the C-terminus of TPL/TPR proteins through a DLNxxP motif. Jsi1 expression in *A. thaliana* leads to the induction of genes related to the ERF-branch of the JA/ET signaling pathway beside inducing also other TPL-target genes. The activation of the JA/ET signaling cascade by *U. maydis* increases biotrophic susceptibility, consequently promoting fungal infection (Darino et al., 2021). Altogether, these findings implicate that TPL/TPR proteins are a hub for *U. maydis* effectors and likely play an important role in the interaction with its host.

In the present study, we performed a systematic screening of 297 putative *U. maydis* effectors for their ability to interact with maize TPL protein RAMOSA 1 ENHANCER LOCUS 2 Like 2 (RELK2) and identified and characterized three novel interactors which we named as TPL interacting protein 6, 7, and 8 (Tip6, Tip7, and Tip8) effectors based on their similarity in TPL interaction with previously identified *U. maydis* effectors Tips1-5 (Bindics et al., 2022). All newly identified Tips induce auxin signalling *in planta* and an octuple Tip mutant, lacking Tip1-8, show virulence defects in comparison to the solopathogenic progenitor strain.

## Results

### Screening and Identification of three novel TPL interacting effectors from *U. maydis*

Recently, we identified seven effector proteins of *U. maydis* which interact at distinct domains with maize TPL proteins and turn on mainly either jasmonate/ethylene or auxin signaling in plants (Darino et al., 2021; Bindics et al., 2022; Navarrete et al., 2022). These studies demonstrate that maize TPL/TPR proteins are a hub for *U. maydis* effectors. To maximize our view on TPL-interacting, auxin signaling inducing *U. maydis* effector candidates we did a two-stepped screen. In step one we screened for TPL interaction whereas in step two we looked at auxin signaling induction by TPL interacting candidates. By yeast two-hybrid (Y2H) screen we tested 297 putative *U. maydis* effector candidates for their capability to interact with maize TPL protein RELK2, a strong interactor for six previously characterized *U. maydis* effectors. For this Y2H screen, RELK2 was fused to the binding domain (BD) of the GAL4 transcription factor, and 297 *U. maydis* putative effector proteins (lacking their signal peptides) were translationally fused one by one to the activation domain of the GAL4 transcription factor (Alcantara et al., 2019). This interaction screen led to the confirmation of the six known RELK2 interacting effectors (Darino et al., 2021; Bindics et al., 2022; Navarrete et al., 2022) and allowed us to identify three additional effector candidates i.e. UMAG_11060, UMAG_05300, and UMAG_05308, that interacted with RELK2 in Y2H assays under low and high stringency conditions (Figure 1A, S1A & B). Additionally, we also tested the interaction of newly identified effector candidates with other maize TPL proteins (REL2, RELK1, and RELK3). Results of this Y2H indicated, that all three effector proteins interact also with RELK3, which is a close homolog of RELK2 (Liu et al., 2019), under high stringency conditions. In contrast to this, all three effectors interacted only under low stringency conditions with RELK1. The weakly expressed REL2 shows only under low stringency condition interaction with UMAG_05300, and UMAG_05308 but not with Umag_11060 (Figure 1A, S1A&B). We named these newly identified TPL/TPR interactors as Tip6 (UMAG_11060), Tip7 (UMAG_05300), Tip8 (UMAG_05308).

**Figure 1:**
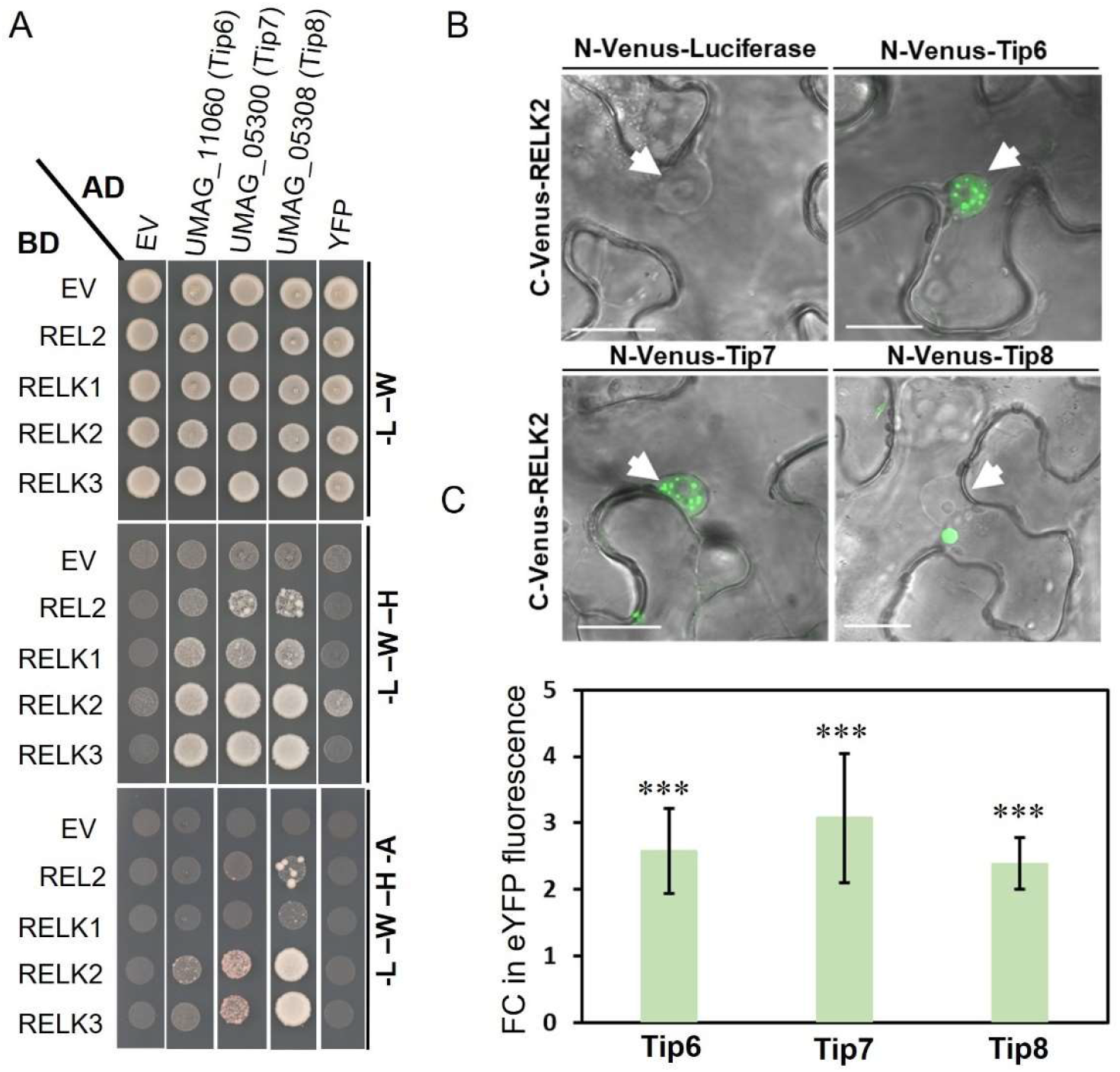
Yeast two-hybrid screening of RELK2 identified three novel auxin signaling inducing maize Topless (TPL) interacting effectors of *Ustilago maydis*. (**A**) Y2H assay showing the interaction of three novel effectors, UMAG_11060 (Tip6), UMAG_05300 (Tip7), UMAG_05308 (Tip8) lacking their signal peptides with maize TPL proteins. Yeast transformants were grown on SD medium lacking the indicated components for auxotrophic selection. Growth on media lacking Leu (L) and Trp (W) was used as a transformation control. Growth on medium or high stringency media lacking L, W, His (H), and Ade (A) indicates protein interactions. (B) Bimolecular fluorescence complementation assays showing the interaction of C-Venus-HA-RELK2 with Tip6, Tip7, and Tip8 (as -Myc-N-Venus tagged) as a fluorescent signal in the *N. benthamiana* epidermal cells but not with the luciferase-Myc-N-Venus control protein. Pictures were taken 2 dpi. The white arrows point to nuclei, Scale bar, 20μm. (C) Fold induction of DR5-eYFP auxin signaling reporter by mCherry tagged Tip6, Tip7 and Tip8 when transiently co-expressed in *N. benthamiana* leaves compared to mCherry as a negative control. Data represent the average of three independent replicates. Error bars indicate standard deviation and asterisks indicate a p-value of less than 0.005 in a student’s t-test.

We also tested the interaction of Tip6, Tip7 and Tip8 with RELK2 *in planta*. For this purpose, a N-terminal part of Venus (Myc-N-venus; Gookin and Assmann, 2014) was attached respectively to Tip6, Tip7 and Tip8 (lacking their signal peptides) and luciferase as a negative control, and a C-terminal part of Venus (HA-C-Venus) was attached to the C-terminal part of RELK2. *A. tumefaciens* carrying *RELK2:HA-C-Venus* plasmid were co-infiltrated into *N. benthamiana* leaves with respective *A. tumefaciens* strains bearing Myc-N-Venus tagged Tip6-8 or luciferase one by one. Confocal Microscopy of transformed leaf epidermal cells was performed two days post infiltration. The bimolecular fluorescence complementation (BiFC) assay results confirmed the interaction of UMAG_11060, UMAG_05300, and UMAG_05308 with that of RELK2 as evident by Venus fluorescent signal in epidermal cells, whereas no fluorescent signal was observed where split Venus RELK2 was co-expressed with split Venus luciferase negative control. Notably, the BiFC signal was observed very specifically for Tip6 / RELK2 and Tip7 /RELK2 interaction in sub-nuclear bodies, whereas Tip8 / RELK2 interaction was detectable in a distinct cytoplasmic, round body next to the nucleus (Figure 1B).

In step two we tested the ability of Tip6, Tip7 and Tip8 to induce auxin signaling *in planta*. For this we transiently co-expressed N-terminally mCherry tagged Tip6, Tip7 and Tip8 lacking their secretion signal (Figure 3A) in the leaves of *N. benthamiana* by *A. tumefaciens*-mediated transformation along with an auxin-responsive *DR5::eYFP* reporter construct (Ulmasov et al., 1997). Three days post infiltration, the fluorescence of the DR5::eYFP reporter was measured from leaf discs taken from infiltrated leaves, and mCherry was used as a negative control. The results of this experiment showed that all three Tips could induce the expression of DR5::eYFP reporter expression to various extents in comparison to the negative control (Figure 1C). To summarize, we identified three novel TPL interacting effectors of *U. maydis* which interacted with TPL/TPR proteins of maize upregulated DR5::eYFP expression in *N. benthamiana* in analogy to previously identified sub-cluster 6a *U. maydis* Tip effectors. Interestingly, a longer expression (5dpi) of Tip8 in *N. benthamiana* epidermal cells led to induction of cell death in our experiments (Figure S1C) implicating possible recognition by *N. benthamiana* plants and induction of effector triggered immune responses.

Previously identified Tips were located genetically clustered on chromosome 6A. We checked the genetic location of Tip6, Tip7 and Tip8. Whereas Tip6 is located on chromosome 11, Tip7 and Tip8 are located in the biggest effector cluster (cluster 19) (Kamper et al., 2006) of *U. maydis* on chromosome 19 (Figure S2A). All three genes are transcriptionally upregulated specifically during the biotrophic phase of *U. maydis*. Expression of Tip6 and Tip7 is peaking at day 2 post infection and stays high till day ten after infection. The expression of Tip8 started to raise at day four, where first galls appear and stayed on till day ten after infection of maize seedlings (Figure S2B).

### Tip6, Tip7 and Tip8 interact with the N-terminal part of Topless proteins

Proteins of the TPL family are multidomain proteins and previously identified TPL interacting *U. maydis* effectors bind either to the N-terminal TPD domain or the C-terminal WD40 domain of TPLs (Darino et al., 2021; Bindics et al., 2022; Navarrete et al., 2022). Therefore, we tested with which domain of maize TPL proteins the newly identified TPL effector candidates interact in the Y2H assay. To this end, truncated versions of the maize TPL proteins containing either the N-terminal TPD domain or C-terminal WD40 domain (Darino et al., 2021; Bindics et al., 2022; Navarrete et al., 2022) were fused to the DNA binding domain (BD) domain of GAL4 and tested for their interactions with UMAG_11060, UMAG_05300, and UMAG_05308 effectors. All three effector proteins interacted only with the N-terminal TPD domain of four maize TPL/TPR proteins (Figure 2, S1C), and no interactions could be observed with the WD40 domains (Figure 2).

**Figure 2:**
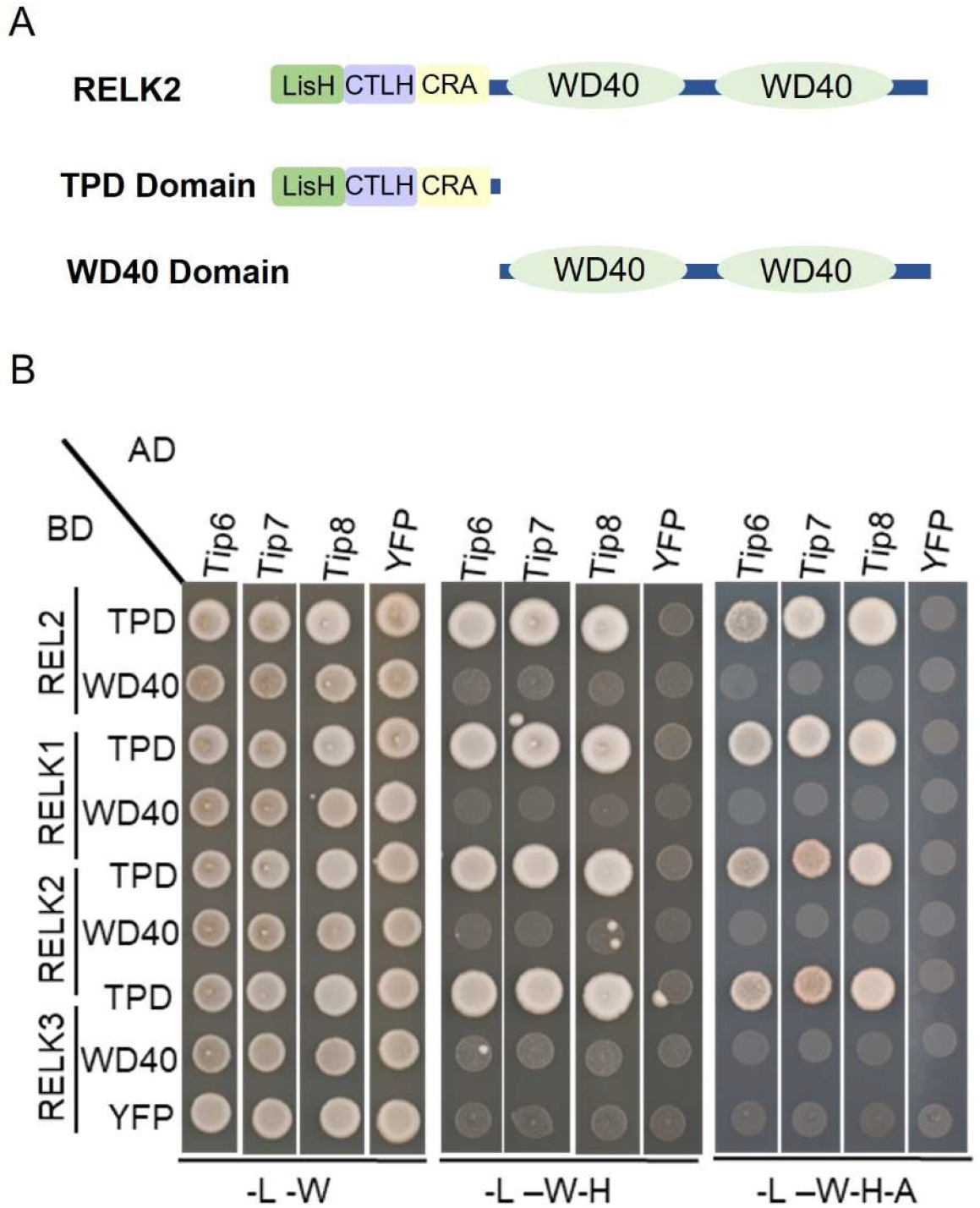
Yeast two-hybrid screening with N- and C-terminal domains of maize RELK2. (A) Schematic presentation of different protein domains of RELK2 (B) Y2H assay showing the interaction of Tip6, Tip7, and Tip8 with C- and N-terminal domain of RELK2 of maize TOPLESS proteins. Yeast transformants were grown on SD medium lacking the indicated components for auxotrophic selection. Growth on media lacking Leu (L) and Trp (W) was used as a transformation control. Growth on medium or high stringency media lacking L, W and, His (H), and Ade (A) indicates protein interactions.

**Figure 3:**
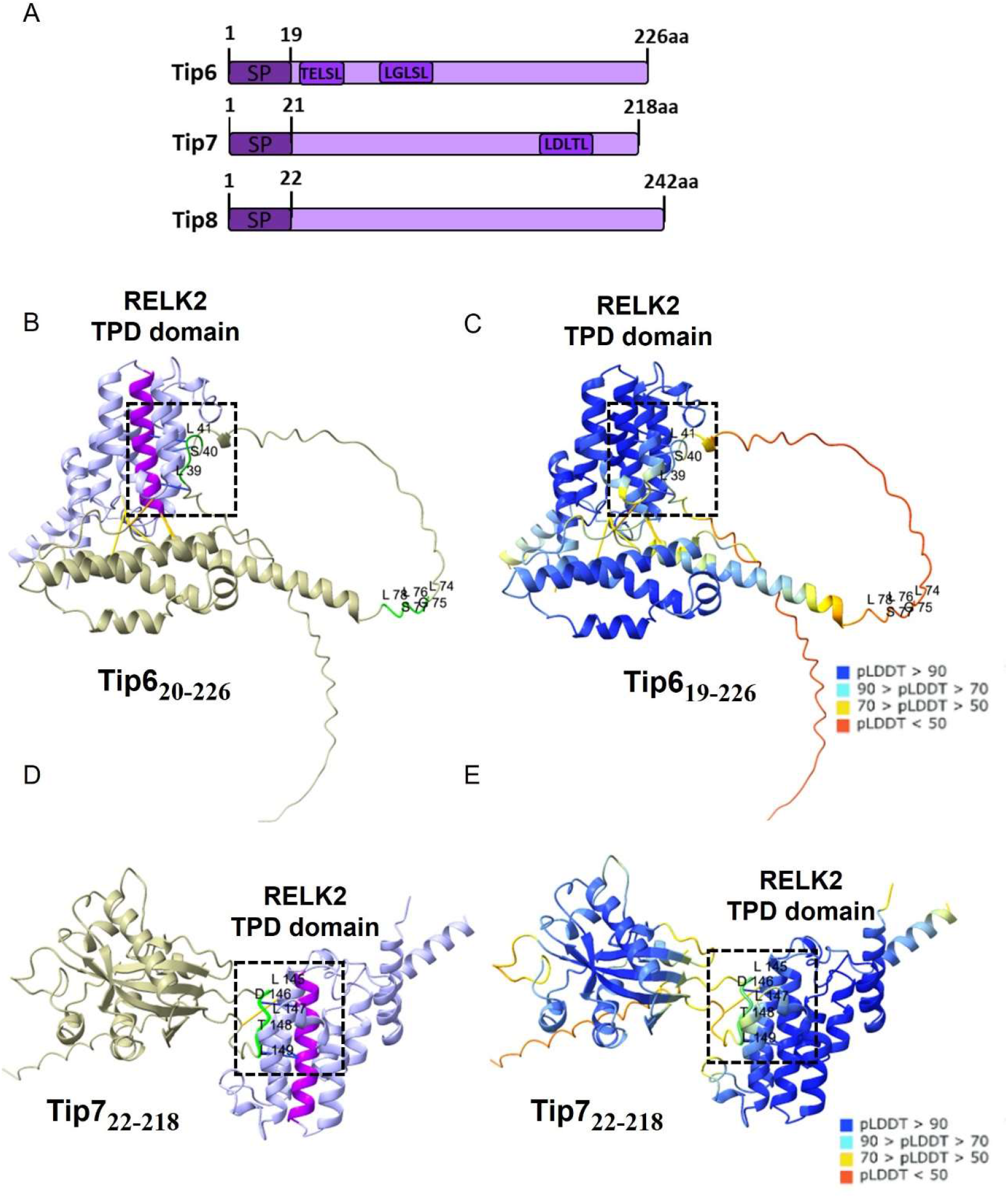
Structure prediction suggest that Tip6 and Tip7 bind with EAR motif to the α helix 5 of the RELK2-TPD domain. **(A)** Schematic representation of full-length Tip effector proteins, the numbers indicate amino acids and SP stands for signal peptide as predicted by SignalP-6.0. (C & E) Alpha fold II structural prediction of Tips and RELK2-TPD domain complex with the color code indicating pLDDT (Local Distance Difference Test) confidence values, blue with a high confidence of above 90 and hence of high accuracy, whereas red colour indicates a lower confidence value. (B & D) False coloring of the model shown in B & D to highlight different motifs and proteins in the complex, beige indicates Tip proteins, green indicates the interaction motifs of Tips to TPD domain, light purple shows TPD domain of RELK2 whereas dark purple indicates the α helix 5 which was shown previously as interaction site of the EAR motif in TPL of *A. thaliana* in crystallographic study (Martin-Arevalillo et al., 2017). The amino acid numbers are counted after removing the signal peptide. The figure was generated with Chimera X with a contact distance of 3 (Goddard et al., 2018; Pettersen et al., 2021), the predicted models are rotated arbitrarily to show clearly the interaction sites.

The TPD domain is composed of three conserved domains: Lisencephaly Homologue (LisH), C-terminal to LisH (CTLH) and CT11-RanBPM (CRA) (Figure 3a). The TPD domain is not only conserved at the sequence level, but also structurally (Ke et al., 2015; Martin-Arevalillo et al., 2017), and is necessary for interaction with EAR motifs present in the transcriptional repressors (Szemenyei et al., 2008). We looked the protein sequences of Tip6-Tip8 for the presence of EAR repressor motif and found that Tip6 and Tip7 both contain a classical LxLxL type ear motif, whereas Tip8 amino acid sequence lacks any previously identified repressor motif. The crystal structures of TPD domains from Arabidopsis and rice are resolved (Ke et al., 2015; Martin-Arevalillo et al., 2017) and a high confidence pLDDT (Local Distance Difference Test) structure of Tip6 and Tip7 by alpha fold II (Jumper et al., 2021; Pettersen et al., 2021) is also predictable. Therefore, next we made a structure prediction of the complexes Tip6-TPD domain of RELK2 and Tip7-TPD domain of RELK2 by using Chimera X software, after removing the signal peptides of Tips. The result of these structural complex analysis predicted a specific interaction of Tip7 EAR motif, more precisely with Aspartate (D) position 146, threonine (T) at position 148 and Leucine (L) at 149, with amino acids of α helix 5 of the TPD domain of RELK2, which has been previous shown to be the interaction domain for the EAR repressor motif (Ke et al., 2015; Martin-Arevalillo et al., 2017). Whereas for Tip6 a binding between Glutamate (E) at position 38, Serine (S) at position 40 and L at 41 was predicted with the residues in α helix 5 of TPD domain of RELK2. Closely looking this motif, TELSL, looks like a reminiscent of classical LxLxL EAR where the first L is missing. To summarize the results, we show that Tip6, Tip7 and Tip8 binds with the N-terminal TPD domain of maize TPL/TPR proteins and the structural prediction further narrow it down to the α helix 5 of TPD domain with Ear motif (LxLxL) in Tip7 and Ear motif like sequence (TELSL) in Tip6.

### Tip6, Tip7, and Tip8 are secreted proteins

To test if the predicted signal peptides of Tip6, Tip7, and Tip8 are indeed functional, we integrated at the CBX-locus of *Ustilago maydis*, constructs coding either for full-length coding sequences of Tips containing the predicted signal peptide or lacking the respective predicted signal peptides (Figure 3A) C-terminal in frame with mCherry fluorescence tag and under transcriptional control of the strong biotrophy-induced promoter of *cmu1* (Djamei et al., 2011). As progenitor strain for transformation, the solopathogenic strain SG200 was used (Kamper et al., 2006). Seven-day-old maize seedlings were infected with the solopathogenic *U. maydis* strains expressing these constructs and confocal microscopy was performed from maize leaf areas with infection symptoms. Whereas *pcmu1*::ΔSP-Tips-mCherry proteins (lacking their signal peptides) stayed inside the fungal hyphae, the full-length Tip-mCherry fusion proteins (*pcmu1*::Tips-mCherry) were secreted into the biotrophic interphase (mCherry signal around the hyphae) between the fungus and the host cell, indicating their secretion (Figure 4A).

**Figure 4:**
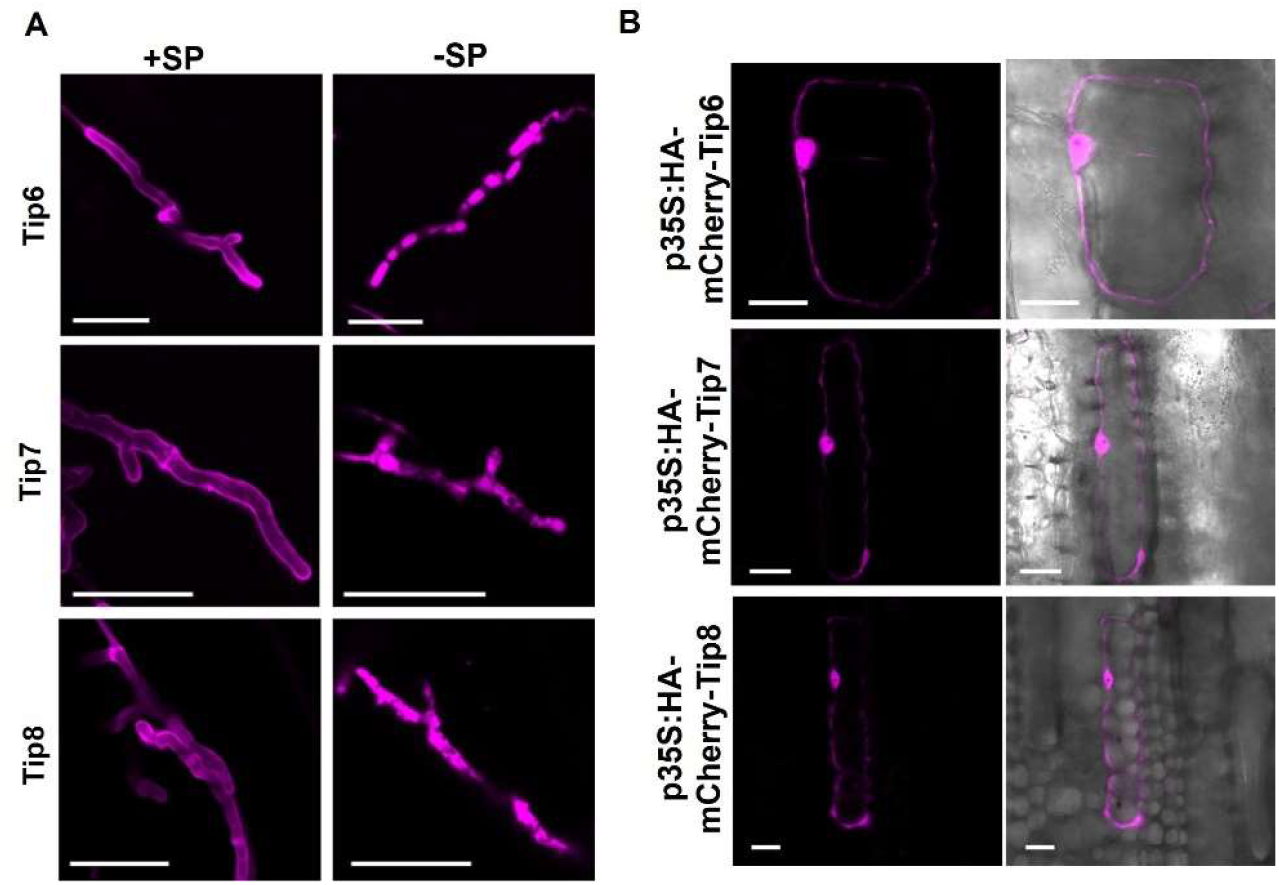
Tip6, Tip7, and Tip8 are secreted effectors that localizes inside the maize cell in the nucleus and cytoplasm. **(A)** Confocal images of maize leaves infected with *U. maydis* strains expressing C-terminally mCherry-tagged Tip6-8 expressed under the c*mu1* promoter with or without signal peptide (SP). Left panel: mCherry signal surrounds the hyphal periphery, indicating secretion into the biotrophic interphase, right panel: mCherry signal retained inside the fungal hyphae, indicating no secretion. Scale bar = 20μm **(B)** Subcellular localization of mCherry-Tip6_20-226_, mCherry-Tip7_22-218_, and mCherry-Tip6_23-242_ in maize leaf epidermis cells after biolistic transformation. For each construct: left panel; mCherry fluorescence only, right panel: merge of bright field and mCherry fluorescence, Scale bar = 20μm.

Then we checked the intracellular localization of Tip effectors in maize host epidermal cells. To this end, we cloned N-terminal mCherry-tagged Tip effectors lacking their signal peptides and introduced these constructs in maize epidermal cells by biolistic transformation. Confocal microscopy reveals a nuclear and cytoplasmic localization as it has been previously also shown for the subcluster 6a Tip effectors (Figure 3B). Taken together, our results show that Tip6, Tip7, and Tip 8 are secreted, fungal proteins and are translocated into the plant cell.

### Secreted Tip6, Tip7 and Tip8 interact in maize infections with TPL proteins and have a role in *U. maydis* virulence

In the next step, we tested if the secreted Tip effectors interact in the native system with TPLs, implicating translocation of the fungal effectors into the host maize nucleus. To perform this, full-length Tip effectors were fused C-terminally to a small hemagglutinin tag (3xHA) under the control of the biotrophy-induced *cmu1* promoter as larger tags like mCherry have been shown to inhibit translocation of *U. maydis* effectors (Tanaka et al., 2015). The mating compatible *U. maydis* strains FB1 and FB2 were transformed with Tip-3xHA expression constructs. As a negative control, FB1 and FB2 strains were transformed with mCherry secreted by the *cmu1* signal peptide and fused to 3xHA tag expression construct. Seven-day-old maize seedlings were infected with a one to one mix of these transgenic FB1 and FB2 strains and five days after infection, leaf material with infection symptoms was collected for protein extraction followed by co-immunoprecipitation (co-IP) with anti-HA antibody. The immunoprecipitates were subjected to western blotting using an anti-TPL antibody (Darino et al., 2021; Bindics et al., 2022; Navarrete et al., 2022). The results showed that all three Tip effectors interacted in the native system with maize TPL proteins as indicated by their ability to pull down TPL proteins in comparison to the negative control secreted mCherry (Figure 5A).

**Figure 5:**
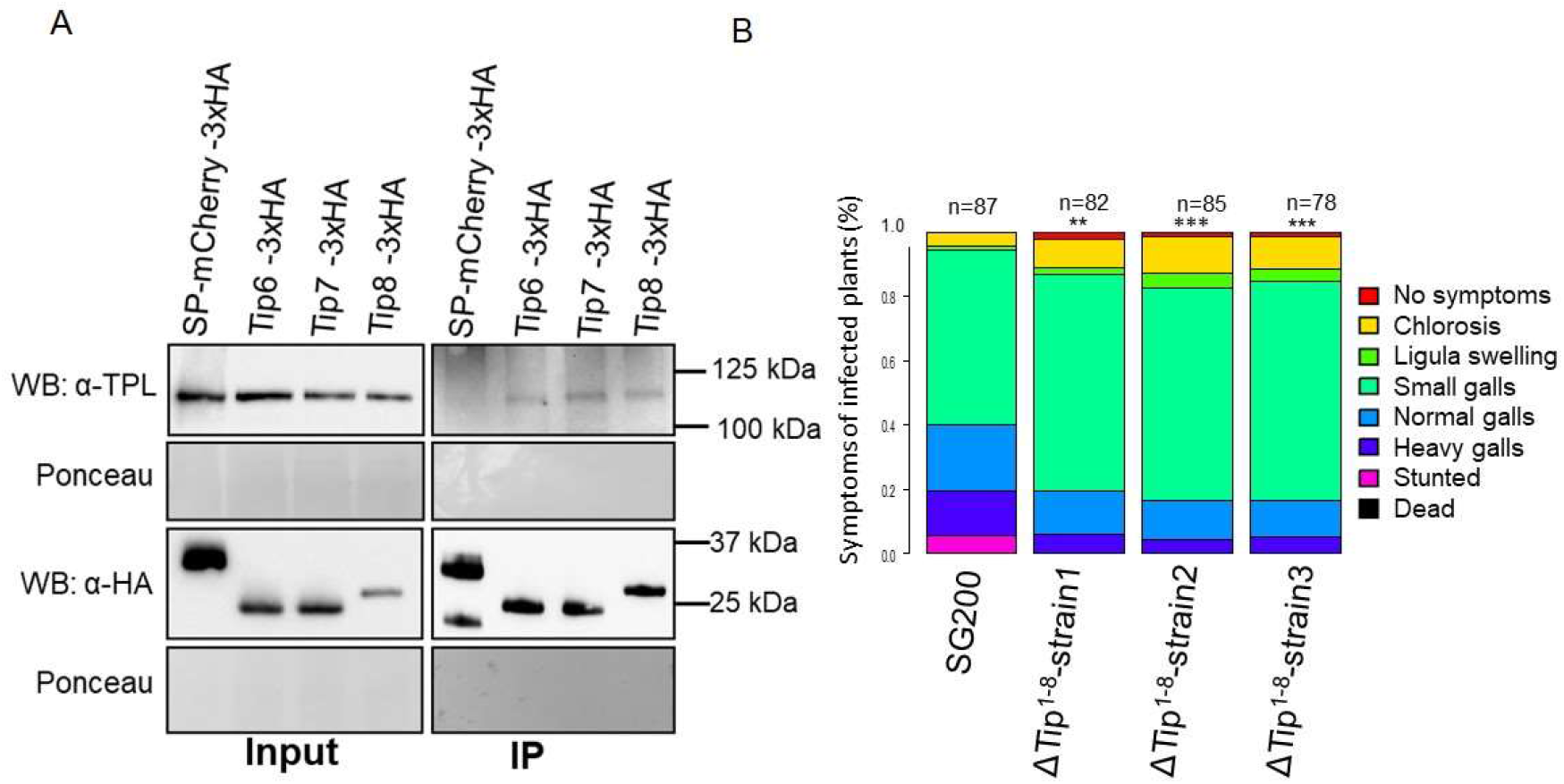
Tip6, Tip7, and Tip8 interact with the maize Topless (TPL) class of proteins after their secretion into the plant cell. **(A)** Protein extracts of maize seedlings infected with *U. maydis* strains secreting Tip6-3xHA, Tip7-3xHA, Tip8-3xHA, or the negative control mCherry-3xHA were subjected to co-immunoprecipitation with an HA-epitope antibody followed by western blotting using either anti-TPL or anti-HA antibodies. Maize TPLs were pulled down specifically with Tip effectors but not from mCherry control. Ponceau staining serves as a loading control. **(B)** *Ustilago maydis* Tips octuple deletion mutants (ΔTip^1-8^) were tested for their ability to infect 7-d-old maize ‘Early Golden Bantam’ seedlings in comparison to SG200. Scoring was performed at 12 d post-infection. Significant differences between strains were analysed by unpaired Wilcoxon test analysis (* *P* < 0.05, ** *P* < 0.01, *** *P* < 0.001)

Next, to test a role of newly identified Tip effectors in virulence, we generated an octuple mutant deletion strains (ΔTip^1-8^) in the solopathogenic strain SG200, where all Tip effectors (Tip1-Tip8) were deleted, by using the CRISPR-Cas9 approach (Schuster et al., 2016; Schuster et al., 2018; Zuo et al., 2020). We evaluated three independent mutants for disease symptoms in *Z. mays* cv Early Golden Bantam. The results of these experiments showed that ΔTip^1-8^ deletion strains are reduced in virulence significantly as compared with the progenitor strain SG200 (Figure 5B).

## Discussion

Previously, we identified and characterized seven effectors of the biotrophic fungal pathogen *U. maydis* targeting proteins of the TPL/TPR family in plants (Darino et al., 2021; Bindics et al., 2022; Navarrete et al., 2022) (Figure 6). To more systematically screen if additional effector candidates target TPL family proteins in plants and induce auxin signaling, we screened 297 effector candidates for their interaction with maize RELK2 and their capability to induce auxin signaling *in planta*. We identified three novel TPL interacting and auxin signaling inducing effectors, Tip6, Tip7, and Tip8 making TPL a central effector hub for *U. maydis*. Previous systematic effector-host protein interaction screens revealed, that convergent evolution drives non-related effectors within a pathogen but also effectors of non-related pathogens to target frequently the same central hubs in the plant immune system (Mukhtar et al., 2011; Wessling et al., 2014). This notion is supported in case of TPL/TPR proteins by the finding, that the oomycete pathogen *Hyaloperonospora arabidopsidis* secretes the HaRxL21 effector that interacts with TPL protein to promote pathogen susceptibility (Harvey et al., 2020). The RolB oncogene protein of rhizogenic Agrobacteria interacts with host plant TPL co-repressors to induce transcriptional changes to promote hairy root development (Gryffroy et al., 2023). *Fusarium oxysporum* effector Six6 interacts with TPL, but gets recognized, triggering SNC1-mediated immune-responses in *A. thaliana* (Gawehns et al., 2014). Similarly, the bacterial *Ralstonia solanacearum* effector PopP2 has an EAR motif being linked to its recognition as an avirulence gene, but also linked to its PTI-suppressing ability as an effector (Segonzac et al., 2017).

**Figure 6:**
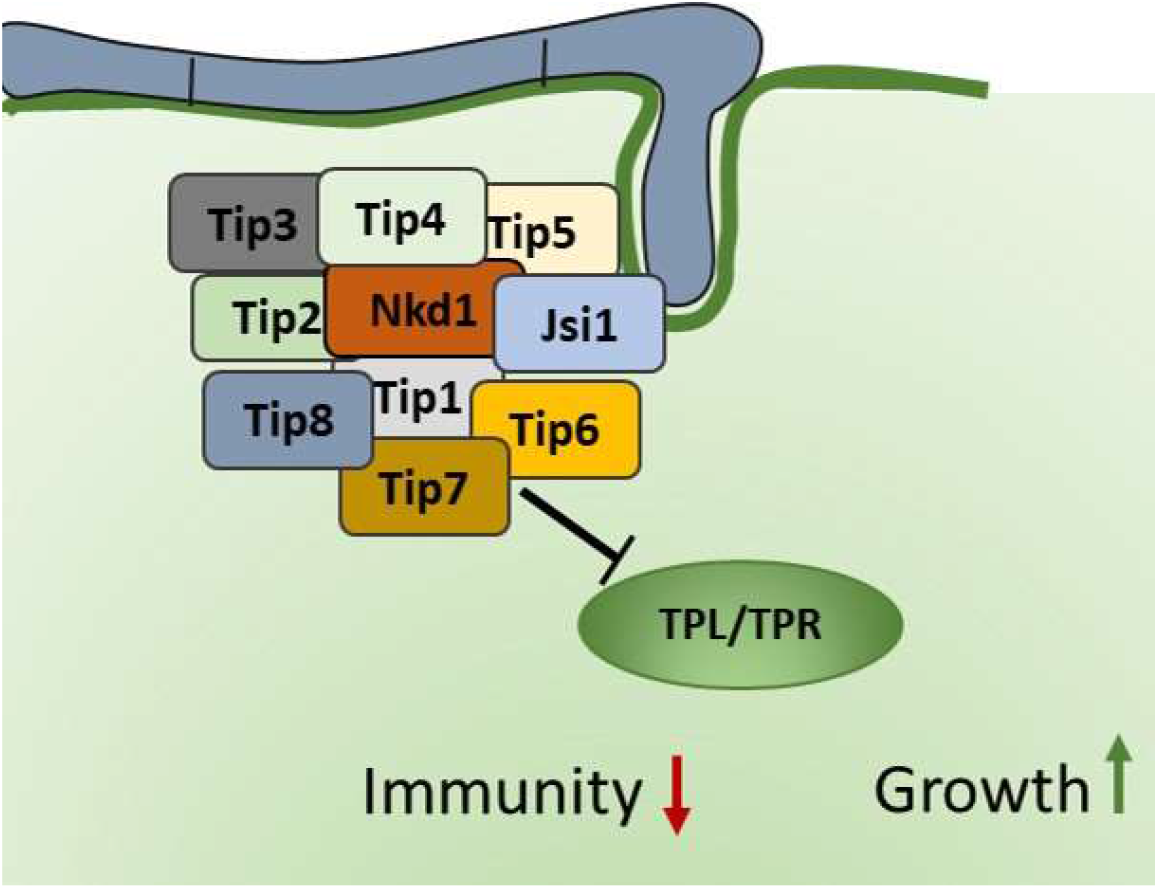
Overview of Topless/Topless related (TPL/TPR) interacting protein effectors of *Ustilago maydis*. The model summarizes the findings of ten TPL/TPR interacting effectors that all negatively modulate TPL/TPR mediated transcriptional repression, thereby inhibiting immune responses (Navarrete et al., 2022) and promote growth signaling pathways (Darino et al., 2021; Bindics et al., 2022; Navarrete et al., 2022).

Whereas Jsi1 is on chromosome 2, Nkd1 on chromosome 5 and Tip1-5 are clustered on chromosome 6a in the genome of *U. maydis*, Tip6 is located on chromosome 11 and Tip7 and Tip8 are both located in the largest effector cluster 19 on chromosome 19. This dispersed distribution of Tips across the genome might be on one hand a result of independent convergent evolutionary events but might also be part of stabilizing forces of functional redundancy and distributed information storage ensuring the important TPL-manipulative functions of the effectome of this biotrophic fungus, even upon genomic redistributions. The generation of the octuple mutant for all known Tip effectors shows a significant reduction in virulence and gall formation.

Tip6 and Tip7 contain classical TPL-binding sites (EAR motifs) similar to Nkd1 effector and Jsi1. The results of our alpha fold II based structural modelling of the Tip7-TPD domain confirmed an interaction of Tip7 with RELK 2 through this EAR motif with high confidence, however, in case of Tip6 the interaction was with the Leucine residue 41 which is located in a motif reminiscent of EAR motif i.e. TELXL, whereas the structural prediction of the part of Tip6 where LxLxL motif is located is with very low pLDDT score and therefore unreliable. On the other side we have Tip8, which lacks any such typical EAR motif in analogy to previously characterized Tips 1-5 effector proteins and it is largely unclear, where and how other six Tip effectors lacking any known TPL binding motifs interfere with TPL /TPR functions. The structural prediction of Tip8 are scoring very low confidence (Figure S3) and thereby it is not possible to predict its binding sites based on the current version from the alpha fold database. All Tip effectors bind to the TPD domain and however for the EAR-motif containing Tip-effectors, their binding to the EAR-binding pocket is not surprising, previous mutant binding studies for Tip1-5 have indicated binding to the amino acid in alpha helix 5, EAR-binding pocket, and thereby influencing the capacity of TPLs to recruiting transcriptional repressors in the plant. As hundreds of plant transcription factors bind to TPL protein family members in Arabidopsis and likely also in all other plants (Causier et al., 2012a), it is understandable why maize did not evolve TPL proteins with alternative binding pockets, as this could only evolve by the extremely unlikely event of parallel mutations in TPL interacting transcriptional repressors in the same individual mutant. Therefore, the interference of *U. maydis* effectors with the EAR-binding pocket in TPL is likely evolutionary favored and convergent evolution not only on the same host target but even on the same host target domain occurred massively in this case. Crystal structure analysis of TPL/TPR proteins (Ke et al., 2015; Martin-Arevalillo et al., 2017) have demonstrated a tetrameric complex of TPL proteins it is an open question if this stoichiometry in terms of Tips in a complex is correlating. Furthermore, it is not clear if different Tips can act simultaneously on a TPL complex and if they act additive for the derepression of certain signaling pathways. The binding predictions as well as mutations in the EAR-binding pocket of TPL followed by interaction assays for Tip1-5 (Bindics et al., 2022) indicate on one hand binding to the same region but not to the same amino acid residues of TPL/TPR proteins. This supports the concept of additive activities of Tips on TPL/TPRs during infection.

The observed cell-death induction due to Tip8 overexpression in *N. benthamiana* is likely a result of an indirect recognition of Tip8 activity on TPL/TPR proteins and resembles plant responses to *Fusarium oxysporum* effector Six6 (Gawehns et al., 2014) or *Ralstonia solanacearum* effector PopP2 (Segonzac et al., 2017). The guarding of TPL/TPRs is likely, as we observe cell-death induction also for several other sequence-wise unrelated Topless-interacting effectors (Figure S1C). As *U. maydis* is a biotroph, its effectome must ensure in sum the suppression of effector triggered immune responses. In the likely case, that maize also guards TPL/TPR activity, a relatively low expression of Tip8 in comparison to other Tips during *U. maydis* biotrophy might be co-evolutionary consequence to prevent massive recognition and immune responses as observed upon overexpression in the heterologous system.

Considering the fact that *U. maydis* is a biotroph, depending on its living host and further the central importance of TPL in manifold developmental and stress-induced processes in the plant, Tips will not simply inactivate all TPL functions but will rather modulate them. Our current working model proposes specific level of binding affinities of Tip effectors leading to interference only with specific subsets of TPL and transcriptional repressor protein complexes, thereby leading to distinct de-repression signatures in the host transcriptome. To gain new insights in the biological processes targeted by Tips and the underlying mechanisms will be an exciting challenge for future research.

## Materials and Methods

### Plant material and plant growth conditions

Maize variety Early Golden Bantam, Old Seeds, Madison, WI, USA was used for assays (virulence, co-immunoprecipitation, localization and secretion assays) and the plants were grown in a temperature-controlled glasshouse (14 h : 10 h, light : dark cycle, 28°C : 20°C). *N. benthamiana* used for infiltrations were grown in controlled short-day conditions (8h light/16h dark at 21°C ± 2°C).

### Accession numbers

RELK1 (Zm00001d040279), RELK2 (Zm00001d028481), RELK3 (Zm00001d047897), REL2 (Zm00001d024523), Tip6 (UMAG_11060), Tip7 (UMAG_05300), Tip7 (UMAG_05308).

### Maize infection assays

*U. maydis* wild-type FB1, FB2 (Kamper et al., 2006) and the solopathogenic strain SG200 and its derivatives were used to infect seven-day-old maize seedlings as described in detail previously (Redkar and Doehlemann, 2016) and symptom scoring was performed at 12 dpi according to (Kamper et al., 2006). Data were analyzed by the Wilcoxon test analysis in R, as described by (Stirnberg and Djamei, 2016).

### Confocal microscopy

Confocal microscopy was performed with a Leica SP8 confocal microscope. mCherry was excited at 561nm and emission was collected between 578-648 nm. Images were processed using the LAS-X software from Leica.

### Molecular cloning

All clonings were performed using green gate cloning system (Lampropoulos et al., 2013). *E. coli* Mach1 (Thermo Fisher Scientific, Waltham, MS, USA) was used for all DNA manipulations and were grown in a dYT liquid medium or on YT agar plates with the required antibiotic supplements. To visualize effector secretion *in planta*, the coding sequences of effectors with or without their predicted secretion signals (Figure 3A) were fused with mCherry at their C-terminus and expressed under the strong biotrophy-induced *cmu1* promoter (Djamei et al., 2011) cloned in a p123 vector derivative (Navarrete et al., 2021) and integrated into the *ip* locus in *U. maydis* solopathogenic strain SG200 (Kamper et al., 2006) by homologous recombination. For co-immunoprecipitation of TPL-effector complexes from infected maize, the coding sequences of effectors were fused with 3xHA at the C-terminus, under transcriptional control of the cmu1 promoter (Djamei et al., 2011) and integrated into the *ip* locus of *U. maydis* mating-compatible strains FB1 and FB2. Secreted mCherry (*pcmu1:spcmu1-mCherry:3xHA*) expressing FB1 and FB2 strains (Bindics et al., 2022) were used as negative control. Deletion mutants of Tip6-8 were created by CRISPR/Cas9 performed as described previously (Bindics et al., 2022). The gRNAs were designed using E-CRISP in the start and end of the genes.

### Co-Immunoprecipitation and transient expression of effectors in maize by biolistic bombardment

Co-immunoprecipitation of Tip5, Tip6, and Tip7 and maize TPLs were performed as described previously (Bindics et al., 2022) by using anti-MYC μMACS™ Micro-Beads (Miltenyi Biotech, Bergisch Gladbach, Germany) according to the manufacturer’s protocol. The input samples and co-IP eluates were detected with anti-TPL (Darino et al., 2021) and anti-HA antibodies. Experiments were repeated at least two times. The biolistic bombardment was performed according to (Bindics et al., 2022).

### Yeast transformations and Y2H assays for TPL interacting effectors

All yeast work was performed according to the yeast protocols handbook (Clontech, Mountainview, CA). For Y2H screening, RELK2 was fused to the Gal4 binding domain in the pGBKT7 vector and transformed into yeast strains AH109 by the LiAc/PEG method. A library of 297 putative effectors *of U. maydis* fused to the Gal4 activation domain transformed into the Y187 yeast strain (Alcantara et al., 2019) was used for mating with RELK2-AD containing AH109 strain. Mated yeasts were plated on SD-W-T double selection and the colonies obtained after three days were further used for spotting assays where the growth of diploids in intermediate or high stringency media five days post-inoculation indicated positive interactions. Positive interactions were verified by the independent transformations of plasmids. For REL2, RELK2, RELK3, and RELK4 Y2H plasmids and Y2H screening was performed as described (Bindics et al., 2022).

## Acknowledgments

The research leading to these results received funding from the European Research Council under the European Union’s Seventh Framework Programme ERC-2013-STG, Grant Agreement: 335691, the Austrian Science Fund (FWF): [P27818-B22, I 3033-B22], the Austrian Academy of Sciences (OEAW), and the Deutsche Forschungsgemeinschaft (DFG, German Research Foundation) under Germany’s Excellence Strategy EXC-2070-390732324 and DFG grant (DJ_64_5-1).

## Author contributions

Conceptualization: AD. Methodology: MK, AD. Investigation: MK, SU, BK, JB, KDI, Project administration: MK. Resources: MK, SU, NN, AD. Original Draft: MK, AD. Funding acquisition: AD, Supervision AD, MK

## Declaration of interests

The authors declare no competing interests

## Notes

### Competing Interest Statement

The authors have declared no competing interest.

