## Supplementary data for "Tip of the iceberg? Three novel TOPLESS interacting effectors of the gall-inducing fungus *Ustilago maydis*"

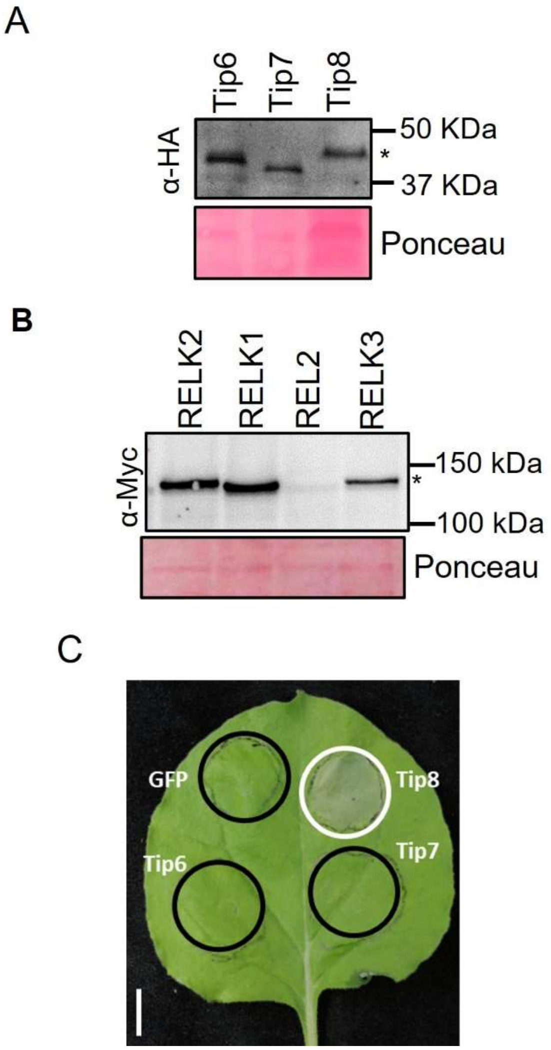


**Supplementary Figure 1. Western blot analysis of proteins from Y2H** (A) Western blot analysis of total protein extracts of yeast used in Y2H and expressing Tips effectors, membranes were incubated with an α-HA antibody. (B) Western blot analysis of total protein extracts of yeast used in Y2H assays and expressing Topless proteins, membranes were incubated with α-Myc antibody. (C) *N. benthamiana* leaf expressing Tip6-8 showing cell-death induction in the case of Tip8 but not for Tip6 and Tip7.

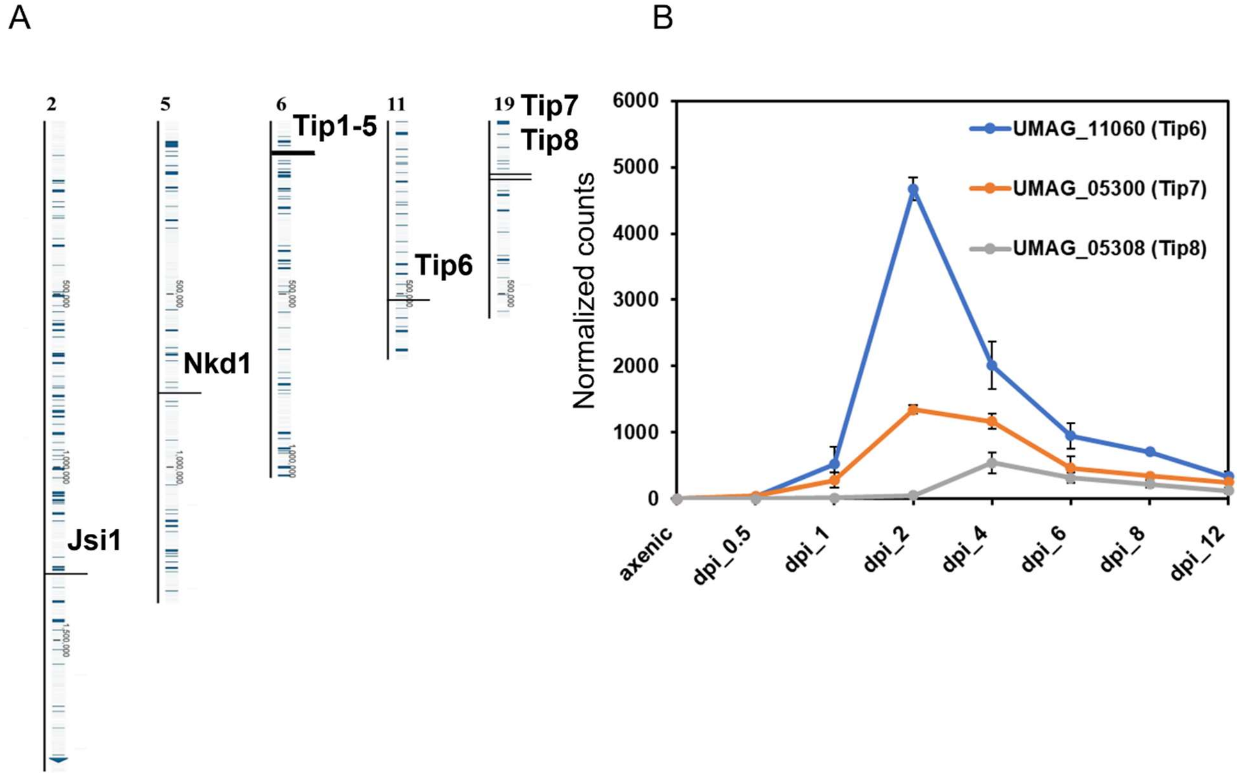


**Supplementary Figure 2. Topless (TPL)- interacting effectors are distributed on different chromosomes and are transcriptionally induced during biotrophy** (A) Chromosomal positioning of TPL- interacting effectors. The numbers above the chromosome indicate the chromosome on which Topless-interacting effectors are located. (B) The graph presents the transcript abundance of Tip6, Tip7, and Tip8 effectors over the course of infection in maize seedlings. Data extracted from (Lanver et al., 2018). Each time point indicates three independent biological replicates. Error bars indicate standard deviation and dpi stands for days post-infection.

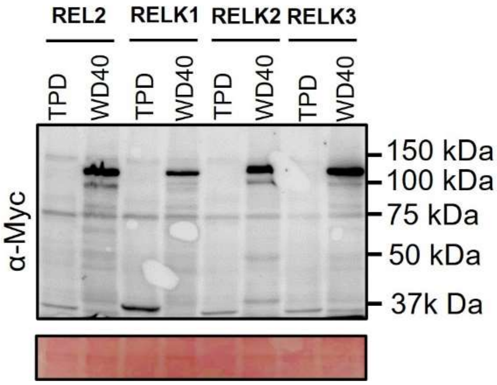


**Supplementary Figure 3. Western blot analysis of proteins from Y2H .** Western blot analysis of total protein extracts of yeast used in Y2H assays and expressing truncated versions of maize Topless proteins, membranes were incubated with α-Myc antibody. Ponceau staining shows loading.

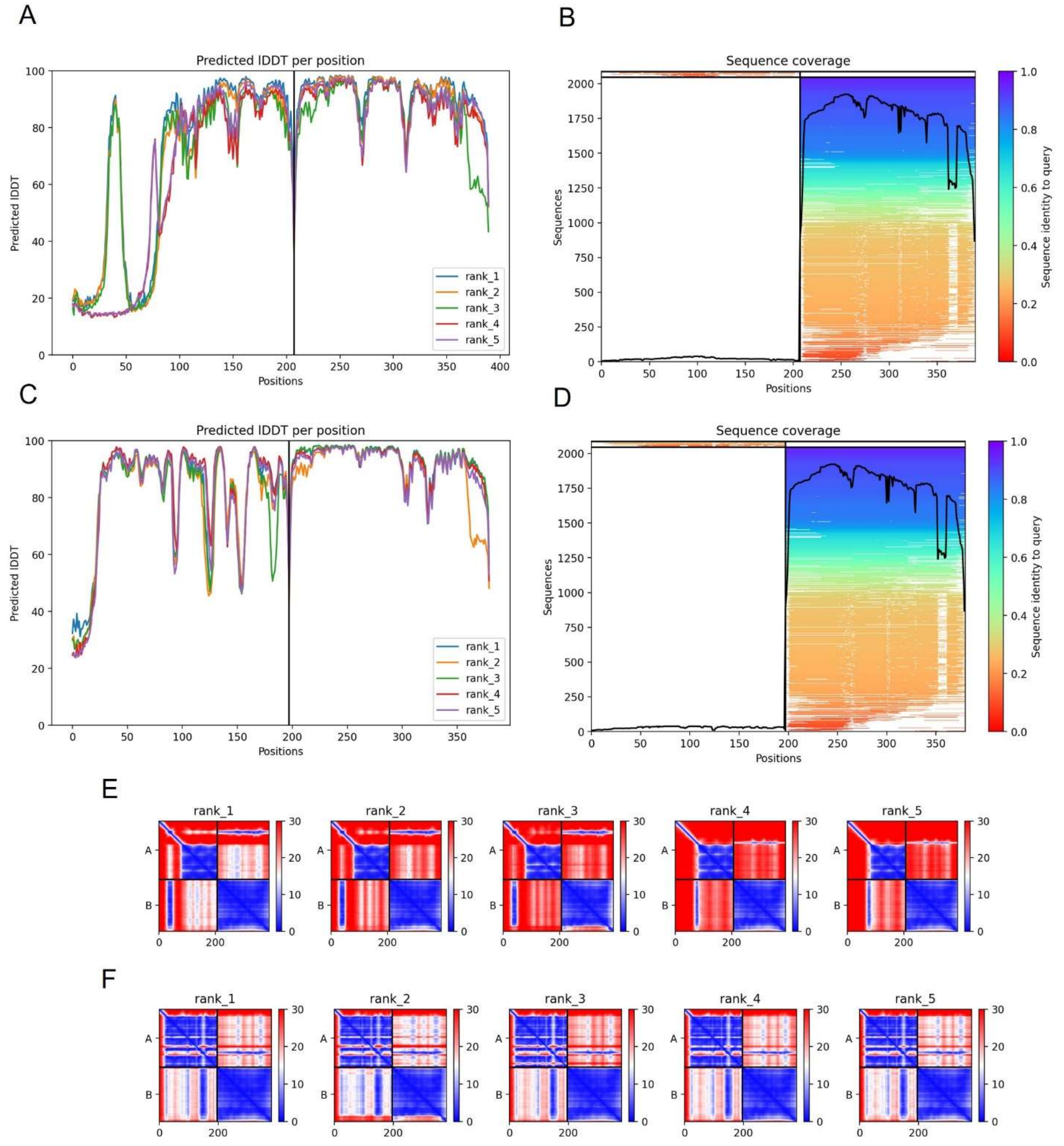


**Supplementary Figure 4. Confidence metrics for the predicted structure of Tip6 and Tip7 complex with TPD domain of RELK2.** (A & C) Predicted local distance difference test (LDDT) score per position for the five models generated by alphafold2 for Tip6 and Tip7 complex with TPD domain of RELK2 respectively. The amino acid position is plotted against the predicted LDDT. pLDDT Values above 90 indicate very high accuracy, values between 70 and 90 indicate a high accuracy, and Values between 50 to 70 indicate a lower accuracy, but it is likely that the predictions of individual secondary structures are correct. (B&E) Sequence coverage for Tip6 and Tip7 complex with TPD domain of RELK2 respectively. (E& F) Prediction aligned error (PAE) score for models. The uncertainty in the predicted distance of two amino acids is color-coded from blue (0 Å) to red (30 Å), as shown in the right bars.

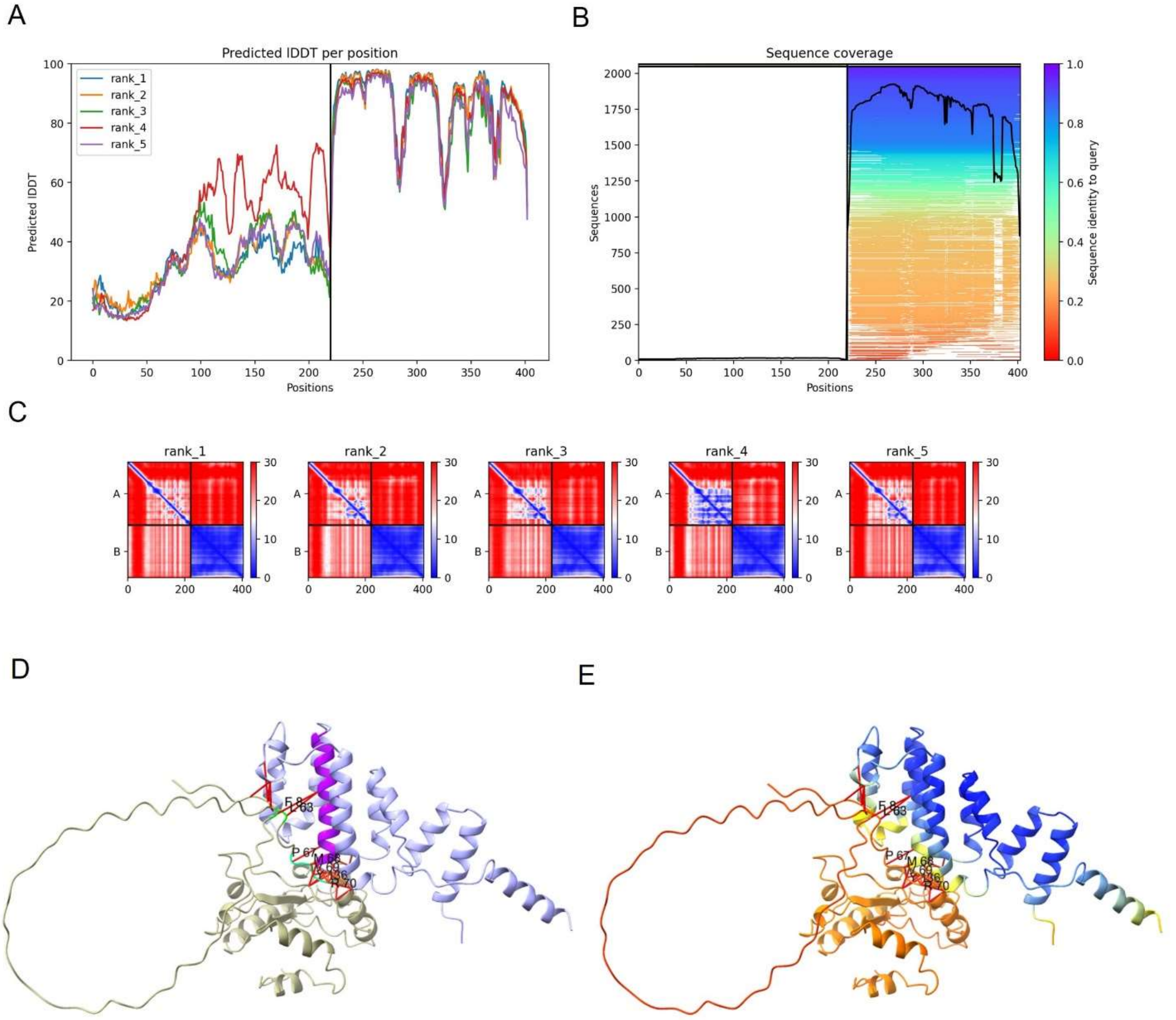


**Supplementary Figure 5. Predicted structural model and confidence metrics for the predicted structure of Tip8 complex with the TPD domain of RELK2.** (A) Predicted local distance difference test (LDDT) score per position for the five models generated by alpha fold II for Tip8 complex with TPD domain of RELK2 respectively. The amino acid position is plotted against the predicted LDDT. pLDDT Values above 90 indicate very high accuracy, values between 70 and 90 indicate a high accuracy, and Values between 50 to 70 indicate a lower accuracy, but it is likely that the predictions of individual secondary structures are correct. (B) Sequence coverage for Tip8 complex with the TPD domain of RELK2 respectively. (C) Prediction aligned error (PAE) score for models. The uncertainty in the predicted distance of two amino acids is color-coded from blue (0 Å) to red (30 Å), as shown in the right bars. (D) False coloring of the model shown in B &

E to highlight different motifs and proteins in the complex, beige indicates Tip proteins, green indicates the interaction motifs of Tips to TPD domain, light purple shows TPD domain of RELK2 whereas dark purple indicates the α helix 5 which was shown previously as interaction site of the EAR motif in TPL of *A. thaliana* in the crystallographic study (Martin-Arevalillo et al., 2017). (E) Alpha fold II structural prediction of Tip8 and RELK2-TPD domain complex with the color code indicating pLDDT confidence values, blue with a high confidence of above 90 and hence of high accuracy, whereas red colour indicates a lower confidence value.
